# Assessing the specificity of the relationship between brain alpha oscillations and tonic pain

**DOI:** 10.1101/787283

**Authors:** Elia Valentini, Sebastian Halder, Daisy McInnersey, Jason Cooke, Vincenzo Romei

**Affiliations:** University of Essex, Department of Psychology and Centre for Brain Science, Colchester, United Kingdom; University of Essex, School of Computer Science and Electronic Engineering, Colchester, United Kingdom; Marie Curie Palliative Care Research Department, Division of Psychiatry, University College London, United Kingdom; University of Bologna, Department of Psychology, Bologna, Italy

**Author notes:** Correspondence should be addressed to Elia Valentini, Centre for Brain Science, Department of Psychology, University of Essex, Wivenhoe Park, Colchester CO4 3SQ, UK.

**Keywords:** alpha, brain, EEG, IAF, pain, unpleasantness.

## Abstract

Recent research has pointed to alpha brain oscillations as a potential clinical biomarker for sensitivity to pain. In particular, researchers claimed that the slowing of individual alpha frequency (IAF) could be an objective marker of pain during prolonged nociceptive stimulation. However, prolonged pain has been also associated with increased IAF. To date, there is insufficient evidence to conclude that IAF can be a neural marker of pain.

The current study aimed at elucidating the conflicting findings by assessing the specificity of the relationship between brain alpha oscillations and tonic pain. We recorded electroencephalography (EEG) on healthy volunteers during exposure to consecutive 5-minute sessions of painful hot water immersion, innocuous warm water immersion and an aversive, non-painful auditory stimulus, matched by unpleasantness to the painful condition. Participants rated stimulus unpleasantness throughout each condition. We also asked participants to sit still with eyes closed and eyes open right before and after the three experimental conditions in order to obtain a robust estimate of baseline alpha. Our findings revealed both increased and decreased IAF during tonic hot stimulation depending on the alpha range targeted (7-13 Hz vs. 8-10 Hz). In addition, they provide mild evidence for a negative relationship between IAF and the unpleasantness of the experience. Finally, we could not identify a difference between IAF during tonic hot temperature and during tonic auditory experience. Altogether, these findings emphasize a twofold frequency pattern (increase vs. decrease) for IAF during tonic thermal pain, thus indicating the need for robust methodological scrutiny of IAF as a neural marker of pain.

## Introduction

Pain is a major societal burden. In the UK chronic pain is estimated to affect between one-third and one-half of the population. This substantive estimate is likely to increase with the progressive ageing of the population (Fayaz, Croft, Langford, Donaldson, & Jones, 2016). This is a societal challenge which requires converging effort of biomedical, psychological and social sciences.

One of the most outstanding questions is how chronic pain originates and develops within the central nervous system, particularly in the brain. Unfortunately, the study of the relationship between the experience of pain and its neural substrates is a remarkable example of reverse inference in the scientific endeavour, and it has seen consensus moving from the hypothesis of no specialized pain centre in the brain (Melzack, 1990; Melzack & Loeser, 1977) to the belief of a “pain matrix” specifically coding the experience of pain (Apkarian, Bushnell, Treede, & Zubieta, 2005; Ingvar, 1999; Tracey & Mantyh, 2007; Treede, Kenshalo, Gracely, & Jones, 1999) until the acceptance, once again, of the lack of pain-specific brain responses as measured with current neuroimaging technology (Davis, Bushnell, Iannetti, St Lawrence, & Coghill, 2015; Iannetti & Mouraux, 2010; Salomons, Iannetti, Liang, & Wood, 2016). However, the endemic lack of neural specificity in the study of pain did not (after all) discourage the search for markers of modulatory mechanisms involved in pain and similar aversive/unpleasant affective bodily states.

In this respect, the examination of the temporal characteristics of pain gains a central role. Experimental studies mostly set out the provocation of either phasic/transient acute pain (milliseconds to seconds) or tonic pain induced via prolonged stimulation (minutes) (Sinke, Schmidt, Forkmann, & Bingel, 2015). While the latter is conceived to be more relevant for the understanding of the neural mechanisms of chronic pain than the former, a greater deal of research has investigated the effects of brief acute pain than prolonged tonic pain on neural responses (Verne, Robinson, & Price, 2004). Over the years a few neuroimaging studies focused on the neural correlates of tonic pain by means of both hemodynamic and electrophysiological techniques (e.g. Casey, Minoshima, Morrow, & Koeppe, 1996; Huber, Bartling, Pachur, Woikowsky-Biedau, & Lautenbacher, 2006; Schreckenberger et al., 2005). Tonic pain seems to trigger hemodynamic activity similar to that elicited during acute pain but with a significant more sustained recruitment of prefrontal regions (Hashmi et al., 2013). Studies recording electroencephalography (EEG) during prolonged pain classically reported decreased alpha power (Chang, Arendt-Nielsen, Graven-Nielsen, Svensson, & Chen, 2001; Chen & Rappelsberger, 1994; Huber et al., 2006). However, this evidence is also commonly associated with pain-unrelated processes (Ploner, Sorg, & Gross, 2017 for a review). In brief, researchers have yet to reliably identify features of neural oscillations (e.g. magnitude, frequency, and phase) that may represent predictive measures of the experience of pain and its modulation. Nevertheless, the systematic assessment of alpha oscillations, and particularly its frequency, has become a target of hope for pain neuroscientists as already happened in other research domains (Cecere, Rees, & Romei, 2015; Cooke, Poch, Gillmeister, Costantini, & Romei, 2019; Foxe & Snyder, 2011; Klimesch, 1999; Mierau, Klimesch, & Lefebvre, 2017; Migliorati et al., 2019; Samaha & Postle, 2015).

This measure has been recently adopted in the context of neurogenic inflammatory pain model (i.e. intradermal capsaicin) on healthy humans (Furman et al., 2018), an approach that allows to induce hyperalgesia, which is one of the main symptoms in chronic pain conditions (Reichling & Levine, 2009). Importantly, Furman et al. (2018) found that reduced individual alpha frequency (IAF) decreased with the increase of pain sensitivity. Interestingly, the study of oscillatory brain activity in patients with chronic pain shows that they exhibit slowed alpha oscillations compared to healthy controls (de Vries et al., 2013; Lim, Kim, Kim, & Chung, 2016; Sarnthein, Stern, Aufenberg, Rousson, & Jeanmonod, 2006; Walton, Dubois, & Llinás, 2010). These studies lend support to the idea that the slowing of IAF, particularly in its lower range (8–9.5 Hz) may contribute to the generation of clinical and chronic pain.

However, there is also evidence of increased alpha frequency during prolonged (5 min) tonic thermal pain in healthy individuals, and a positive correlation between the increase of alpha frequency and the increase of pain perception (Nir, Sinai, Raz, Sprecher, & Yamitsky, 2010). In their most recent study Furman et al. (2019) suggest that the slowing of IAF correlates with prolonged pain rather than repeated consecutive phasic painful stimulation. Crucially, this correlational finding has been interpreted as evidence of IAF being a reliable biomarker of pain.

Here, we devised a study aimed at investigating the relationship between alpha oscillations and tonic painful somatosensory stimulation. Our main aim was to test whether acceleration or slowing of alpha would single out during tonic pain compared to several control conditions as well as testing the strength of the correlation between IAF and perception. To attain our goal, we recorded EEG on healthy volunteers during exposure to consecutive 5 min sessions of painful hot water immersion, innocuous warm water immersion and an aversive prolonged non-painful auditory stimulation, which aimed to dissect the specificity of alpha frequency for the prolonged hot water immersion. In order to establish a perceptual/experiential compatibility between the two conditions we focused our psychophysical assessment on the unpleasantness of their experience. The rationale being that unpleasantness is the most distinctive feature of pain (Merskey et al., 1979; Price, 2000) while being strongly correlated with the intensity, salience, and the homeostatic threatening value of the somatosensory stimulation (Borsook, Edwards, Elman, Becerra, & Levine, 2013; Price, 2000; Price et al., 2002). During the experiment participants rated stimulus unpleasantness throughout each condition. We also asked participants to sit still with eyes closed and eyes open right before and after the three experimental conditions. Based on previous research, we tested the confirmatory hypotheses that (1) individual alpha power (IAP) is reduced during prolonged hot water immersion compared with innocuous conditions, (2) IAF is slowed in prolonged hot water immersion compared with innocuous conditions, (3) IAF during baselines and hot water immersion is negatively correlated with unpleasantness ratings of a prolonged hot water immersion. We also tested the explorative hypothesis that (4) IAF is slowed during prolonged hot water immersion compared with an equally unpleasant prolonged auditory experience.

To test these hypothesis we developed a twofold approach to the analysis of alpha oscillations. We first run an assumptions-free analysis whereby we used the widest alpha range (7-13 Hz) to perform an unrestrained observation of alpha modulations. As second complementary analysis, we performed an a priori analysis of the lower spectrum of the alpha band (8-12 Hz) according to previous observations that pain conditions are associated with a slowing down of alpha rhythm (within 8-10 Hz).

## Materials and Methods

### Participants

A total of 43 participants volunteered to take part in the study which was approved by the ethics committee of the University of Essex. Seven participants were excluded. One participant disclosed to have had taken a painkiller prior to the experiment. Another participant failed the perceptual matching procedure (see below for detail). Data from the 5 remaining participants were excluded due to technical issues with EEG recording.

This resulted in a final sample size of 36 participants (22 female, mean age: 25.36, age range: 20–56). Twenty-four participants self-identified as White/Caucasian, 6 identified as Asian/Pacific Islander, and 6 as Other. All participants had normal or corrected-to-normal vision and normal hearing. Prior to attending, the study screening took place to ensure that participants had no history of neurological, psychiatric or pain disorders that could interfere with the study or jeopardise their safety.

### Sensory and pain stimulation

We used immersion in hot water to induce a tonic sensation of thermal discomfort. This was induced by immersing the left hand up to the wrist in a 30L tank (RW-3025P, Medline Scientific) with constantly circulating hot water, initially set to 45 °C. This specific temperature has previously been shown to induce a moderate level of pain in healthy subjects (e.g. Granot et al., 2008). Importantly, we instructed our participants to focus on the unpleasantness of their experience as this would have been the dimension across which they would have assessed the other experimental conditions too.

To create a neutral (non-painful) control condition participants immersed their left hand up to the wrist in the same water tank, with the water temperature set to be 6°C lower than the temperature used for the hot condition. This 6°C reduction was selected based on pilot sessions to define the optimal temperature reduction to obtain a minimally unpleasant/no unpleasant water temperature.

Finally, in order to create an unpleasant auditory stimulus with a level of discomfort comparable to that of the hot water, a constant high-frequency tone (5000 Hz, Saw Tooth waveform) was created in Audacity (v. 2.0.5) software and played through a pair of noise-cancelling headphones (NC-40, Lindy). This condition was devised to control for the negative affect contribution to the alpha oscillations. While being not painful, this condition allowed us to exclude that the modulation of alpha activity is not brought about by non-pain-related negative affective states. The volume of the auditory stimulus was determined through the perceptual matching process detailed in the following section.

### Perceptual matching procedure

Anecdotally, some participants reported initial difficulty associated with the qualitative difference between the experience of somatosensory pain and the discomfort associated to the non-painful and non-noxious but distressing sound. Nevertheless, all the participants eventually reported the hot temperature as a painful experience. We informed the participants that the sound would not have had a “painful” quality but that they were required to strive to detect a loudness level that would have generated a similar level of unpleasantness. They were specifically required to focus on the unpleasantness of the sensory stimulation. The goal of the matching procedure was to ensure that the unpleasantness of the auditory stimulus and hot water condition did not significantly differ from one another during the experimental session.

In an exact replica of the conditions during the experiment, participants were seated ∼65cm from a screen, with the water bath placed to their left, and a mouse and volume adjustment knob within easy reach of their right hand. Participants wore noise cancelling headphones throughout the full procedure.

Participants were required to place their left hand up to the wrist into a hot bath (45°C) and were instructed to find a comfortable position and to keep their hand as still as possible. Towels were used for padding to keep any pressure on the participant’s wrist at a minimum (to avoid loss of sensation) and to maximise their level of comfort. Participants were then asked to rate the unpleasantness of their sensation on an onscreen visual-analogue scale (VAS), with verbal anchors at 0 (“No unpleasant”) and 100 (“Intolerable unpleasantness”) and numerical markers at 25, 50 and 75. The scale appeared every 10 seconds for 2.5 minutes. If unpleasantness was consistently rated between 50 and 75, the participant then progressed to the next stage of the matching process. If the participant could not tolerate 45°C, the temperature was reduced by 0.5°C and the matching procedure was started again from the beginning. Similarly, if the participant consistently rated the unpleasantness below 50 on the VAS, the temperature was increased by 0.5°C and the matching procedure started again. This correction process was completed as many times as deemed necessary to ensure that ratings were within the required range. Yet participant’s safety and comfort was maximised.

After approximately 2.5 minutes (15 VAS ratings) at the final selected temperature, participants were offered a short break, before the auditory stimulus began playing through the headphones. Participants were instructed to remain in the water bath whilst this block took place, they were asked to increase the volume of the sound such that the unpleasantness of the auditory stimulus matched the unpleasantness of the sensation elicited by the hot immersion. For the following ∼2.5 minutes, participants rated the affective component associated with the auditory stimulus on the onscreen VAS (using the identical scale from the previous block). Again participants were offered a short break before commencing the final matching block.

In the final block participants were asked to remove their hand from the hot water. Here they only listened to the auditory stimulus in isolation for a further ∼2.5 minutes, whilst continuing to rate the unpleasantness on the onscreen scale (once again using the same scale as before). They were reminded to adjust the volume to maintain a level of unpleasantness equal to magnitude of unpleasantness they recalled during the hot immersion. This further rating phase allowed the participant to compensate potential overestimation associated with the multisensory enhancement occurring in the previous rating task. Prior to commencing the matching block participants were asked to refrain from touching the volume control knob any more during the study once the matching task was completed. This level of volume (and the corresponding water temperature) was then used for that participant for the remainder of the experimental procedure. We eventually subtracted 6 °C from the final temperature of the hot water condition to obtain the target temperature for the warm water condition. This amount was empirically chosen in a preliminary phase whereby several lab members tried several temperature reductions. We came to the conclusion that this was the most effective quantity in compellingly reducing unpleasantness without producing a floor effect on unpleasantness in most of the pilot trials.

### EEG recording

Prior to any measurements, sixty-two Ag/AgCI electrodes (Easycap, BrainProducts GmbH, Gilching, Germany) were mounted to obtain EEG recordings (Synamps RT, Neuroscan, Compumedics). The electrodes were carefully placed according to the 10-20 International System. The impedance of all electrodes was kept below 10 kΩ, and the EEG signal was amplified and digitised at 1000 Hz. The online reference was placed upon the left earlobe and the ground was located at electrode position AFz. A secondary offline reference was also used, this was placed upon the participant’s right earlobe.

### Experimental design and procedure

Fig.1 depicts the experimental design and procedure. Each participant was submitted to five experimental conditions: ‘eyes closed*’*, ‘eyes open’, ‘hot’, ‘warm’, ‘sound’. Closed and open eyes conditions were baseline EEG measurements requiring no sensory rating, whereas the three tonic sensory conditions required ratings of unpleasantness. Prior to the perceptual matching task, the EEG cap was mounted upon the scalp, and the signal was quality-checked. No recording took place during the matching task. During the experimental task the EEG was recorded in seven separate blocks corresponding to each separate baseline (eyes-open and eyes closed) and sensory condition (hot, warm, sound). Eyes closed and eyes open were recorded both at the beginning and at the end of the experimental session (their trial order was counterbalanced across participants). More specifically, participants rested with their eyes closed (or open) for 2.5 minutes and then with their eyes open (or closed) for a further 2.5 minutes.

**Figure 1.**
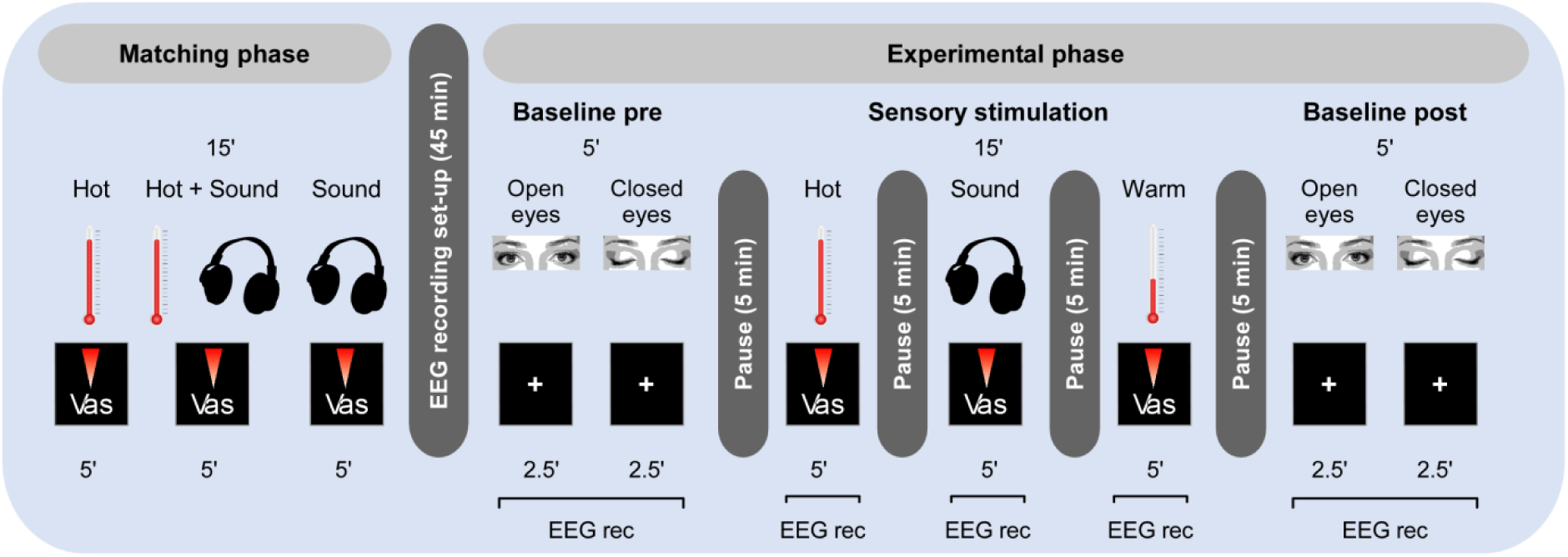
Procedure and design. Participants were comfortably seated with a water bath placed to their left, and a mouse and volume adjustment knob within easy reach of their right hand. Participants wore a pair of noise cancelling headphones throughout (see main text for a full description). They first underwent a matching procedure to determine the water temperature and volume level that induced a target unpleasantness rating (range: 50-75) on a visuo-analogue scale (VAS) ranging between 0 (“not unpleasant”) and 100 (“intolerable unpleasantness”). The resulting parameters were then used in the experimental phase, during which the EEG was recorded in different blocks. The figure shows one of the possible combinations of events. The order of the different blocks was counterbalanced across participants. Baseline recordings (pre and post) consisted of two 2.5-minute blocks (eyes open or closed). A fixation cross was shown on screen while participants were simply asked to sit in a comfortable position and relax. After a 5-minute break, participants completed the tonic Hot, Sound and Warm blocks. Throughout these three blocks, participants rated the unpleasantness of the sensation every 15-seconds on the same VAS as in the matching phase.

Participants were then presented with the 3 types of sensory conditions (i.e. tonic hot, tonic warm, tonic sound), which were counterbalanced across participants so as to remove as much influence of order effects as possible; each time being asked to rate the level of unpleasantness on the VAS every 10 seconds. During these trials (and the open eyes baseline condition) a central fixation cross was presented to participants in order to hold their attention. Participants were first submitted to the tonic hot condition, for example, for ∼5 minutes (30 VAS ratings), then to the tonic warm condition for another ∼5 minutes and finally to the tonic sound condition for a final ∼5 minutes. Finally, a post procedure baseline measurement was once again recorded. As before, participants rested again with their eyes open (or closed) for 2.5 minutes and closed (or open) for a further 2.5 minutes. Between each block participants were offered a 5-minute resting period.

#### Data analysis

##### Psychophysics

Normality was assessed for all the data. Perceptual matching ratings did not violate normality and therefore were analysed using T-test. Mean unpleasantness ratings during the main experiment were averaged for each participant across tonic hot, warm and sound conditions and submitted to non-parametric repeated-measures one-way Friedman ANOVA with Holms corrected pairwise comparison were run to assess differences in dependent variables between conditions. Cohen’s d are reported as measures of effect size.

##### EEG data-analysis – pre-processing

The continuous EEG data were pre-processed with EEGLAB – an open-source tool-box run within the MATLAB environment (Delorme & Makeig, 2004). Individual participants’ data were first re-sampled at 500 Hz and then band-pass filtered from 0.9 to 100 Hz (filter order 8). We removed DC and used Cleanline to remove power line-related sinusoidal artefacts (50 and 100 Hz) (http://www.nitrc.org/projects/cleanline). Data were then re-referenced to the average of the earlobes and submitted to the Artefact Subspace Reconstruction (ASR) plugin (Mullen et al., 2013). We used only the “burst” and “window” criteria with conservative values of 20 and 0.5 respectively. These criteria allowed us to keep all the stimulus events within each recording block while correcting the signal from significant artefacts before further processing them using independent component analysis (ICA, (Makeig, Bell, Jung, & Sejnowski, 1996) to subtract eye and muscle-related artefacts, aided by the multiple artefact rejection algorithm (MARA, Winkler, Haufe, & Tangermann, 2011). Data resulting from the ICA were cropped in equally sized data chunks (i.e. 320s each).

##### Identification of alpha oscillatory activity and analysis of its power and frequency

To isolate the power and frequency of the alpha rhythm we followed the procedure outlined by Furman et al., 2018. We first applied a band-pass filter from 5 to 16 Hz. We used principle component analysis (PCA) to restrict the number of components examined to the number of components remained after initial MARA-based artefact subtraction. We then ran the extended version of Infomax ICA on the number of independent components (ICs) available in the sample (mean: 32.44, SD: 5.51, range: 20-43). Next we visually inspected topographies, time courses and power spectra of the extracted components and selected those components with a peak in the alpha range that was not modulated by the eyes closed condition. We then stored an unmixing matrix that only included the selected components. Each experimental condition was filtered using this unmixing matrix and segmented into regular non-overlapping epochs of 5 seconds length. We calculated the power-spectral density (PSD) by transforming each of these epochs in the frequency domain using a multitaper frequency transformation (Hanning windows, 2-40 Hz, 0.2 Hz bin width) via the use of Fieldtrip (Oostenveld, Fries, Maris, & Schoffelen, 2011). Finally we calculated the center of gravity (CoG; Klimesch, Schimke, & Pfurtscheller, 1993) of the alpha peak in a 7-13 Hz and an 8-10 Hz window of interest using the following equation:

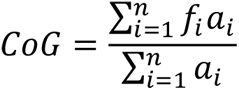

with *n* being the number of frequency bins in the window of interest, *f_i_* being the frequency and *a_i_* the amplitude represented by the *i*^th^ bin. When we selected more than one component with central alpha this calculation was independently performed for each component and then averaged over the frequency and amplitude of the individual components. It is worth noting that a series of different approaches have been developed to assess alpha frequency and extract a summary index of it (e.g. Grandy et al., 2013; Lodder & van Putten, 2011). However, by extracting a weighted mean of the selected frequency range, CoG is meant to reduce the impact of non-canonical individual alpha distributions such as split or multiple peaks (Chiang, Rennie, Robinson, Albada, & Kerr, 2011).

##### Statistical analyses of alpha oscillatory activity

We first examined the relationship between the pre and post alpha baselines using correlation coefficients, then we analysed the relationship between ratings of unpleasantness and both alpha power and frequency in their respective condition as well as their relationship with baseline alpha. As Shapiro-Wilk test revealed CoG alpha power being non-normally distributed (max W=0.55; p_s_<0.001) we used Spearman Rho as correlation coefficient for both dependent variables (DVs). Previous research showed a relationship between alpha oscillations during rest no-sensory setting and pain ratings collected during subsequent painful stimulation (Furman et al., 2018). Hence, we correlated baseline alpha oscillatory activity with unpleasantness ratings obtained in the three different sensory conditions (warm, hot, sound) whereas alpha obtained in each sensory condition was correlated with their respective unpleasantness rating. Similarly, group differences were analysed by a one-way non-parametric analysis of variance (ANOVA) with five levels: eyes closed, eyes open, hot, warm, sound. When the Friedman ANOVA revealed a main effect of the factor ‘condition’ we then calculated Conover’s post-hoc tests to dissect differences between the five levels of the DV. Post-hoc alpha values were corrected using Holm’s approach.

Bayesian Kendall correlations were also computed to test the specific alternative hypothesis that CoG alpha frequency during closed and open eyes as well as during hot stimulation would negatively correlate with unpleasantness ratings reported during hot stimulations, as suggested by recent studies (Furman et al., 2018; Furman et al., 2019). All the analyses were performed using JASP (*JASP Team*, 2018).

## Results

### Perceptual matching

Mean (±SD) temperature (°C) and sound pressure level (dB) resulting from the matching procedure and chosen for the experiment were 44.5 (±0.49) and 90.84 (±11.41), respectively. One participant’s unpleasantness ratings were lost due to a technical software fault. Final ratings for the unpleasant auditory and hot conditions did not deviate from normality (W=0.97; p=0.39). Both distributions were in the desired range as the average ratings were beyond VAS 50 of unpleasantness (hot: 66.87±6.99; sound: 69.59±9.77). Paired t-tests confirmed that the participants successfully achieved a matched experience of unpleasantness for the hot and sound condition (t_34_=-1.62; p=0.11; d=-0.27) before starting the experiment.

### Unpleasantness ratings

As ratings during tonic warm condition were highly skewed (W=0.62; p<0.001) we computed non-parametric ANOVA. The Friedman test revealed a significant difference in unpleasantness between conditions (Chi_2_=54.50; p<0.001). Fig. 2 displays the distribution of average VAS unpleasantness ratings for each condition and the difference between conditions as revealed by the post-hoc tests. Participants rated both tonic hot (67.83±9.18) and sound (64.27±11.93) as similarly unpleasant (T_70_=0.71; p=0.48; d=0.32), thus indicating the matching procedure was successful. As expected, they were both (T_70_=6.72 and T_70_=6.01; p_s_<0.001; d=6.05 and d=4.73) rated as significantly more unpleasant than the tonic warm condition (4.58±6.94).

**Figure 2.**
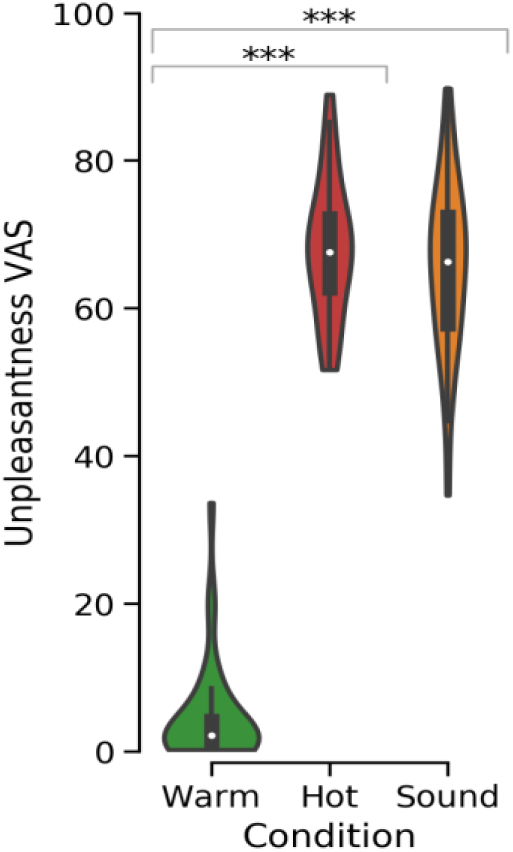
Unpleasantness ratings. Violin-plots representing the median of individual average unpleasantness ratings in the three sensory conditions (x axis). The boxes represent the inner quartiles while whiskers represent data within 1.5 times the inner quartile range. The outer shape is a Kernel density estimation of showing the distribution density of the ratings. Asterisks represent statistical two-tailed significance (***p<0.001). Note that the tonic hot and sound conditions do not differ in unpleasantness while both differ from the somatosensory control condition (i.e. tonic warm).

### Wide spectrum analysis (7-13 Hz)

Fig. 3 shows grand-average ICs PSD profiles (panel A), box-plots illustrating both their CoG power and frequency profiles across the five conditions (panel B), and scalp topographies displaying the mean power at the frequency of the CoG computed from the signal using only the back-projection of the selected alpha ICs.

**Figure 3.**
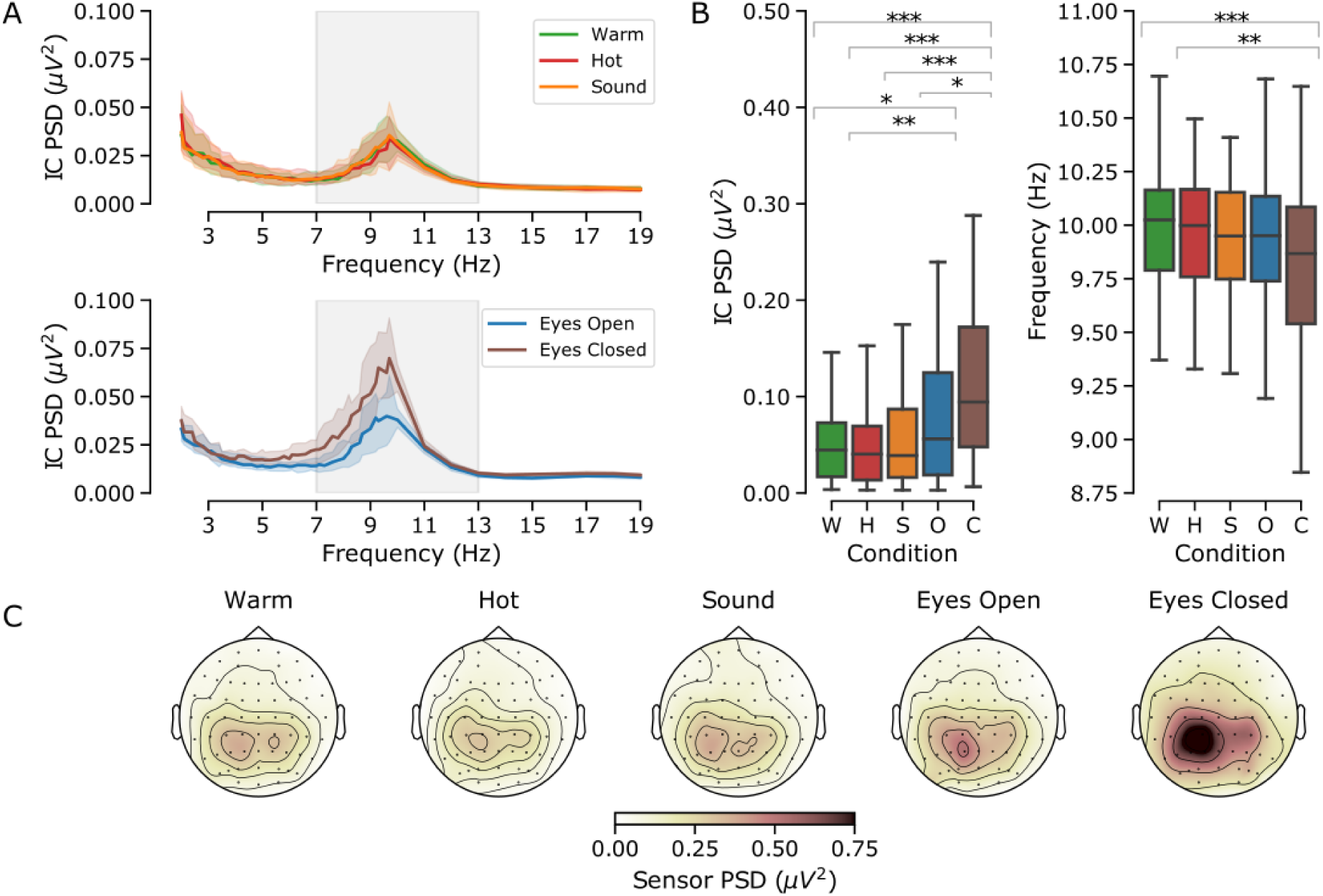
Wide spectrum analysis (7-13 Hz). Panel A: grand-median (shaded area: 95% confidence interval) PSDs obtained following the IC alpha identification phase. Note how the alpha oscillations in the sensory stimulation conditions (upper graph) show lower magnitude and narrower frequency range than the baseline conditions (lower graph). Panel B: box-plots representing the median of individual average CoG power (left graph) and frequency (right graph) in all the five conditions (Warm, W, green; Hot, H, red; Sound, S, orange, Open eyes, O, blue, Closed eyes, C, brown). The boxes represent the inner quartiles while whiskers represent data within 1.5 times the inner quartile range. Data outside this range was not included in this figure. Asterisks represent statistical two-tailed significance (***p<0.001, **p<0.01, *p<0.05). Note the expected progressive reduction of alpha magnitude from the baselines to the sensory stimulation conditions. Note also the increase of frequency in the somatosensory conditions compared with the eyes closed baseline. Panel C: scalp topographies of mean power in the selected CoG frequency based only on the projection of the selected alpha ICs into sensors space.

### Preliminary analysis of the 7-13 Hz baseline alpha

Correlation analysis revealed a strong positive correlation between the first and last recordings of open and closed eyes conditions (see Fig. 1 for a schematic representation of the experimental design) both for the CoG power (Rho=0.91; p<0.001 for both conditions) and frequency (Rho=0.86; p<0.001 and Rho=0.91; p<0.001). We therefore proceeded to average the initial and final session recordings to obtain a single measure of alpha oscillations in the open and closed eyes conditions while accounting for temporal order effects on alpha oscillatory patterns.

### Analysis of the 7-13 Hz alpha power

The difference in alpha magnitude depicted in Fig. 3 is supported by a significant variation of CoG alpha power across conditions (Chi_4_=66.16; p<0.001). This is explained by greater power during eyes closed compared with all the other conditions, and greater power during eyes open compared with tonic warm and hot conditions (Fig 3, panel B, left). The alpha magnitude reduction was not different across the three sensory conditions. These comparisons are summarised in Table 1.

**Table 1.** Summary of post-hoc comparisons for the CoG alpha power as extracted from the 7-13 Hz wide frequency window. Asterisks represent statistical two-tailed significance (***p<0.001, **p<0.01, *p<0.05).

|  |  |  | T-Stat | p value | p<sub>holm</sub> | Cohen's d |
| --- | --- | --- | --- | --- | --- | --- |
| **Eyes closed** | Vs. | Eyes open | 3.19 | 0.002 | 0.01* | 0.32 |
|  | Vs. | Warm | 6.27 | <0.001 | <0.001*** | 0.71 |
|  | Vs. | Hot | 7.05 | <0.001 | <0.001*** | 0.97 |
|  | Vs. | Sound | 5.57 | <0.001 | <0.001*** | 0.52 |
| **Eyes open** | Vs. | Warm | 3.08 | 0.002 | 0.01* | 0.58 |
|  | Vs. | Hot | 3.86 | <0.001 | 0.001** | 0.40 |
|  | Vs. | Sound | 2.37 | 0.02 | 0.08 | 0.25 |
| **Warm** | Vs. | Hot | 0.78 | 0.44 | 0.87 | 0.21 |
|  | Vs. | Sound | 0.70 | 0.48 | 0.87 | 0.03 |
| **Hot** | **Vs.** | **Sound** | 1.48 | 0.14 | 0.42 | -0.10 |

### Relationship between ratings of unpleasantness and 7-13 Hz alpha power

Individual CoG alpha power did not correlate with unpleasantness ratings in any of the planned correlations (Table 2).

**Table 2.**
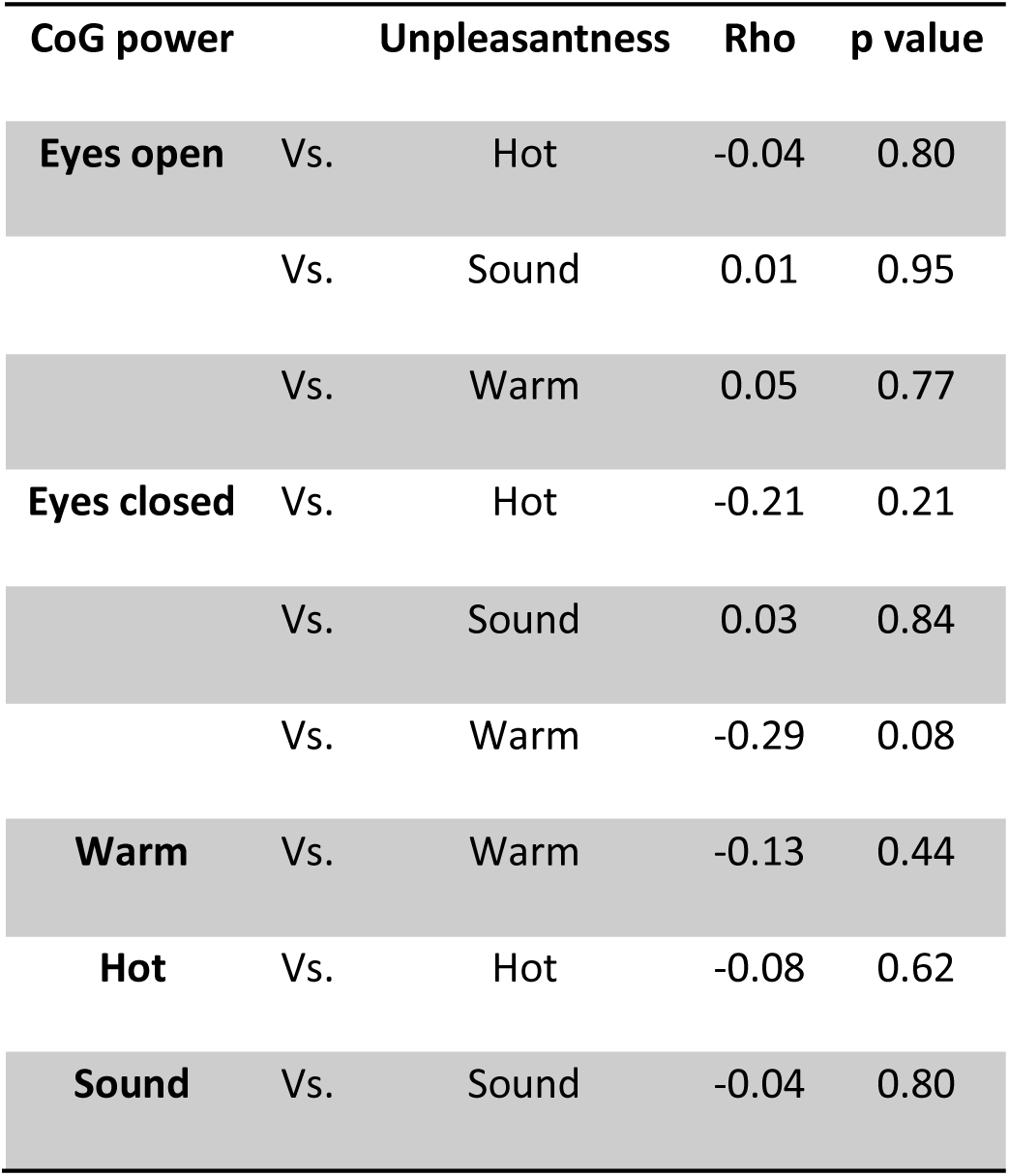
Coefficients of correlation between the CoG 7-13 Hz alpha power and unpleasantness ratings.

### Analysis of the 7-13 Hz alpha frequency

CoG alpha frequency differed across conditions (Chi_4_=24.39; p<0.001). Fig. 3 (panels A and B) shows how this difference was explained by slower frequency during eyes closed compared with the warm and hot condition. However, the alpha CoG frequency was not different across the three sensory conditions. These comparisons are summarized in Table 3.

**Table 3.** Summary of post-hoc comparisons for the CoG alpha frequency as extracted from the 7-13 Hz wide frequency window. Asterisks represent statistical two-tailed significance (***p<0.001, **p<0.01, *p<0.05).

|  |  |  | T-Stat | p value | p<sub>holm</sub> | Cohen's d |
| --- | --- | --- | --- | --- | --- | --- |
| **Eyes closed** | Vs. | Eyes open | 2.30 | 0.02 | 0.16 | -0.68 |
|  | Vs. | Warm | 4.41 | <0.001 | <0.001*** | -0.77 |
|  | Vs. | Hot | 3.82 | <0.001 | 0.002** | -0.69 |
|  | Vs. | Sound | 1.89 | 0.06 | 0.28 | -0.51 |
| **Eyes open** | Vs. | Warm | 2.11 | 0.04 | 0.22 | -0.29 |
|  | Vs. | Hot | 1.52 | 0.13 | 0.39 | -0.22 |
|  | Vs. | Sound | 0.41 | 0.68 | 1.00 | 0.06 |
| **Warm** | Vs. | Hot | 0.59 | 0.55 | 1.00 | 0.15 |
|  | Vs. | Sound | 2.52 | 0.01 | 0.10 | 0.43 |
| **Hot** | Vs. | Sound | 1.93 | 0.06 | 0.28 | 0.43 |

### Relationship between ratings of unpleasantness and 7-13 Hz alpha frequency

Individual CoG alpha frequency did not correlate with unpleasantness ratings in any of the planned correlations (Table 4).

**Table 4.** Coefficients of correlation between the 7-13 Hz CoG alpha frequency and unpleasantness ratings.

| CoG frequency | Unpleasantness | Rho | p value |
| --- | --- | --- | --- |
| **Eyes open** | Vs. Hot | 0.05 | 0.79 |
|  | Vs. Sound | -0.09 | 0.59 |
|  | Vs. Warm | 0.05 | 0.76 |
| **Eyes closed** | Vs. Hot | 0.01 | 0.97 |
|  | Vs. Sound | -0.23 | 0.18 |
|  | Vs. Warm | -0.01 | 0.94 |
| **Warm** | Vs. Warm | -0.01 | 0.95 |
| **Hot** | Vs. Hot | -0.04 | 0.82 |
| **Sound** | Vs. Sound | -0.11 | 0.52 |

### Slow alpha analysis (8-10 Hz)

Fig. 4 shows grand-average ICs PSD profiles (panel A), box-plots illustrating both their CoG power and frequency profiles across the five conditions (panel B), and scalp topographies displaying the mean power computed from the original signal prior to the ICs selection phase (panel C).

**Figure 4.**
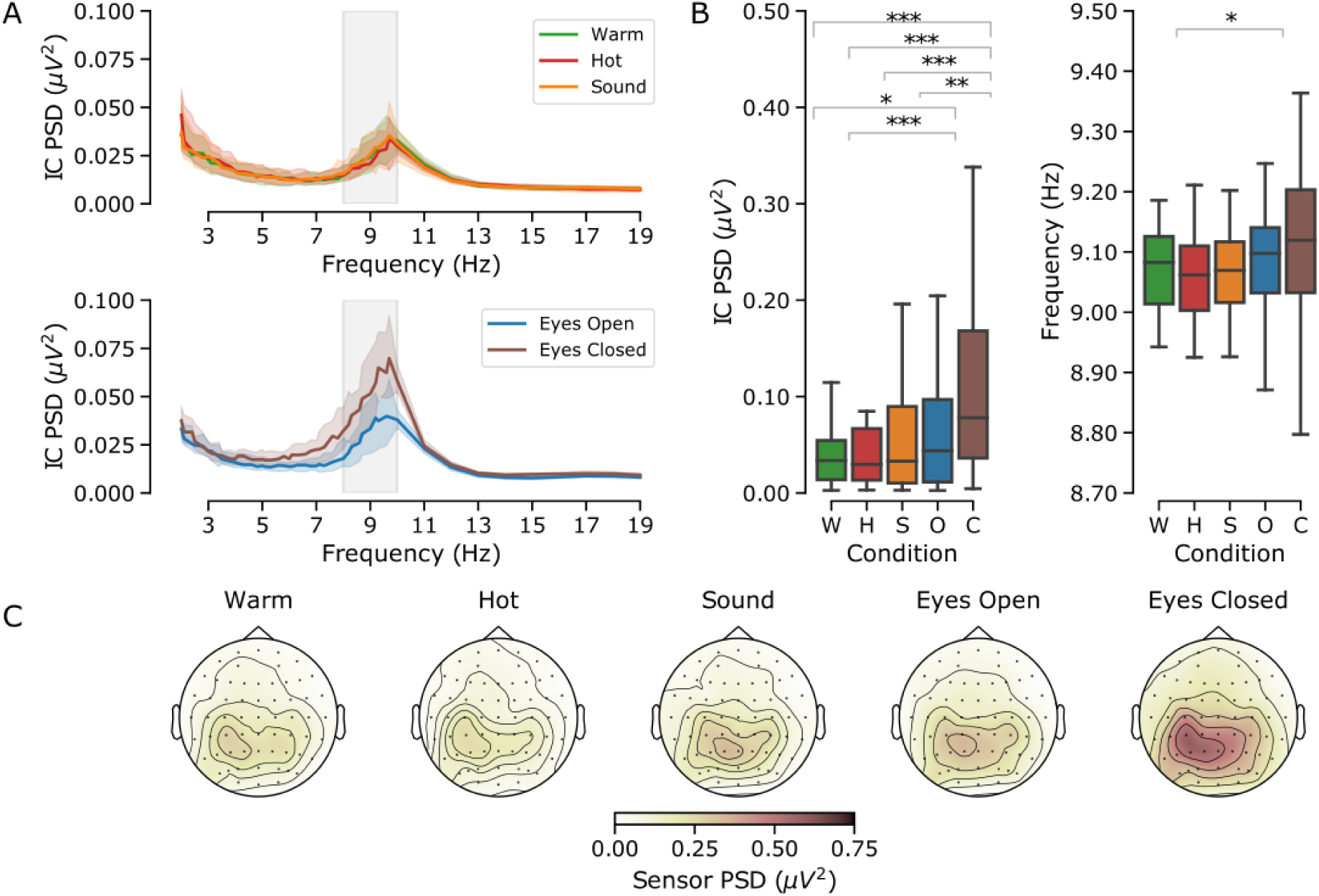
Slow alpha analysis (8-10 Hz). Panel A: grand-median (shaded area: 95% confidence interval) PSDs obtained following the IC alpha identification phase. Note how the alpha oscillations in the sensory stimulation conditions (upper graph) show lower magnitude and narrower frequency range than the baseline conditions (lower graph). Panel B: box-plots representing the median of individual average CoG power (left graph) and frequency (right graph) in all the five conditions (Warm, W, green; Hot, H, red; Sound, S, orange, Open eyes, O, blue, Closed eyes, C, brown The boxes represent the inner quartiles while whiskers represent data within 1.5 times the inner quartile range. Data outside this range was not included in this figure. Asterisks represent statistical two-tailed significance (***p<0.001, **p<0.01, *p<0.05). Note the expected progressive reduction of alpha magnitude from the baselines to the sensory stimulation conditions. Note also the reduction of frequency in the tonic hot condition compared with the eyes open baseline. Panel C: scalp topographies of mean power in the selected CoG frequency based only on the projection of the selected alpha ICs into sensors space.

### Preliminary analysis of the 8-10 Hz baseline alpha

Correlation analysis revealed a strong positive correlation between the first and last recordings of open and closed eyes conditions both for the CoG power (Rho=0.93; p<0.001 and Rho=0.91; p<0.001) and frequency (Rho=0.75; p<0.001 and Rho=0.91; p<0.001). We therefore proceeded to average the initial and final session recordings.

### Analysis of the 8-10 Hz alpha power

The difference in alpha magnitude depicted in Fig. 4 is supported by a significant variation of CoG alpha power across conditions (Chi_4_=70.25; p<0.001). This is explained by greater power during eyes closed and open compared with all the other conditions (Fig 3, panel B, left). The alpha magnitude reduction was not different across the three sensory conditions. These comparisons are summarised in Table 5.

**Table 5.**
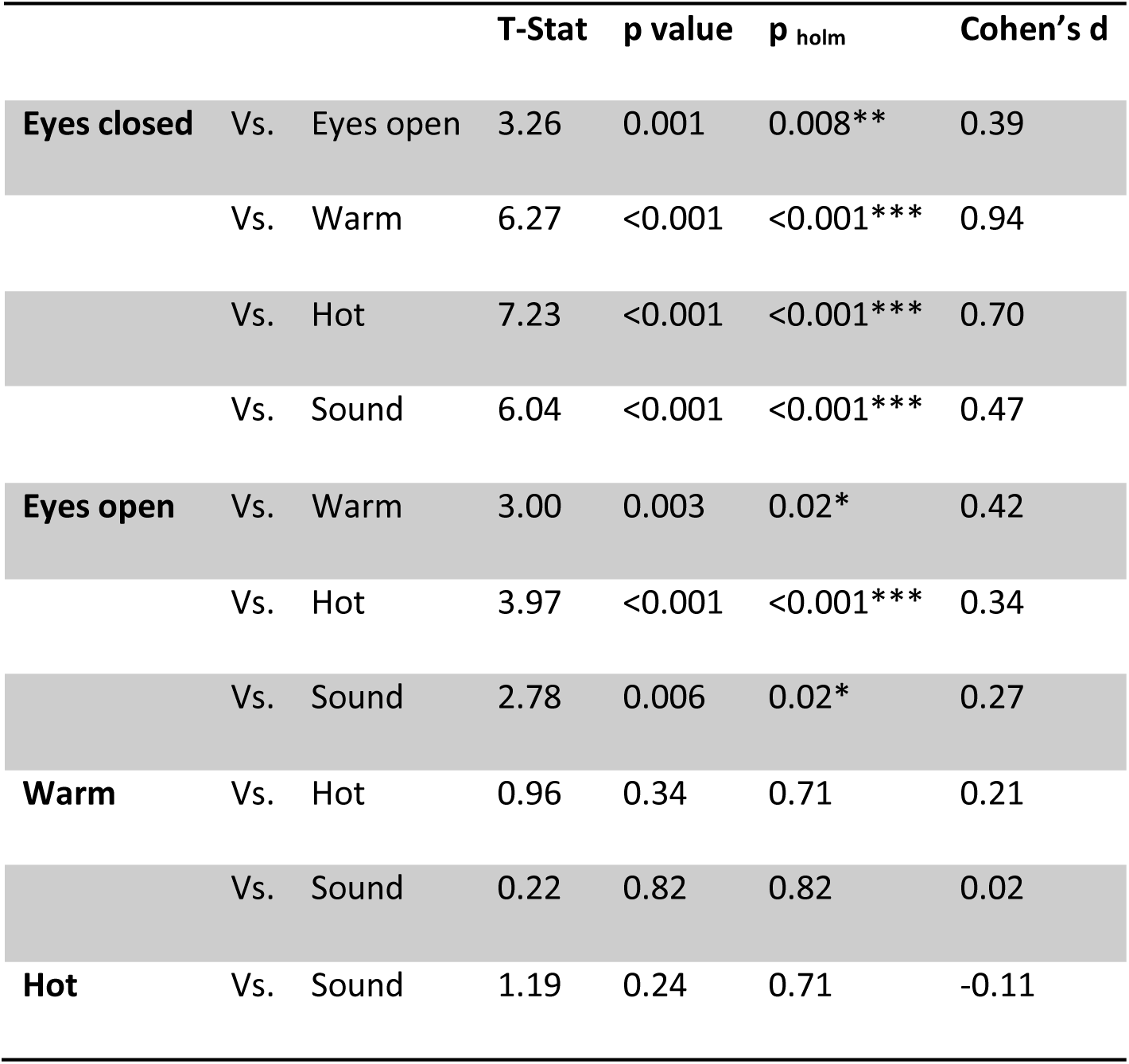
Summary of post-hoc comparisons for the CoG alpha power as extracted from the 8-10 Hz frequency window. Asterisks represent statistical two-tailed significance (***p<0.001, **p<0.01, *p<0.05).

### Relationship between ratings of unpleasantness and 8-10 Hz CoG alpha power

Individual CoG alpha power did not correlate with unpleasantness ratings in any of the planned correlations (Table 6).

**Table 6.** Coefficients of correlation between the 8-10 Hz CoG alpha power and unpleasantness ratings.

| CoG power |  | Unpleasantness | Rho | p value |
| --- | --- | --- | --- | --- |
| Eyes open | Vs. | Hot | -0.12 | 0.47 |
|  | Vs. | Sound | -0.01 | 0.93 |
|  | Vs. | Warm | 0.02 | 0.89 |
| Eyes closed | Vs. | Hot | -0.22 | 0.19 |
|  | Vs. | Sound | 0.05 | 0.79 |
|  | Vs. | Warm | -0.18 | 0.28 |
| Warm | Vs. | Warm | 0.02 | 0.92 |
| Hot | Vs. | Hot | -0.11 | 0.53 |
| Sound | Vs. | Sound | -0.05 | 0.75 |

### Analysis of the 8-10 Hz alpha frequency

CoG Alpha frequency differed across conditions (Chi_4_=10.13; p=0.04). Fig. 4 (panels A and B) shows how this difference was explained by slower frequency during the hot condition compared with eyes open (Fig. 4, panel B, right). However, the alpha CoG frequency was not different across the three sensory conditions. These comparisons are summarised in Table 7.

**Table 7.** Summary of post-hoc comparisons for the CoG alpha frequency as extracted from the 8-10 Hz frequency window. Asterisks represent statistical two-tailed significance (***p<0.001, **p<0.01, *p<0.05).

|  |  |  | T-Stat | p value | p<sub>holm</sub> | Cohen's d |
| --- | --- | --- | --- | --- | --- | --- |
| Eyes closed | Vs. | Eyes open | 0.67 | 0.51 | 1.00 | 0.39 |
|  | Vs. | Warm | 0.78 | 0.44 | 1.00 | 0.29 |
|  | Vs. | Hot | 2.26 | 0.02 | 0.23 | 0.39 |
|  | Vs. | Sound | 1.15 | 0.25 | 1.00 | 0.32 |
| **Eyes open** | Vs. | Warm | 1.45 | 0.15 | 0.98 | 0.27 |
|  | Vs. | Hot | 2.93 | 0.004 | 0.04* | 0.48 |
|  | Vs. | Sound | 1.82 | 0.07 | 0.57 | 0.31 |
| **Warm** | Vs. | Hot | 1.48 | 0.14 | 0.98 | -0.29 |
|  | Vs. | Sound | 0.37 | 0.71 | 1.00 | 0.02 |
| **Hot** | Vs. | Sound | 1.11 | 0.27 | 1.00 | -0.30 |

### Relationship between ratings of unpleasantness and 8-10 Hz alpha frequency

Individual CoG alpha frequency did not correlate with unpleasantness ratings in any of the planned correlations (Table 8).

**Table 8.**
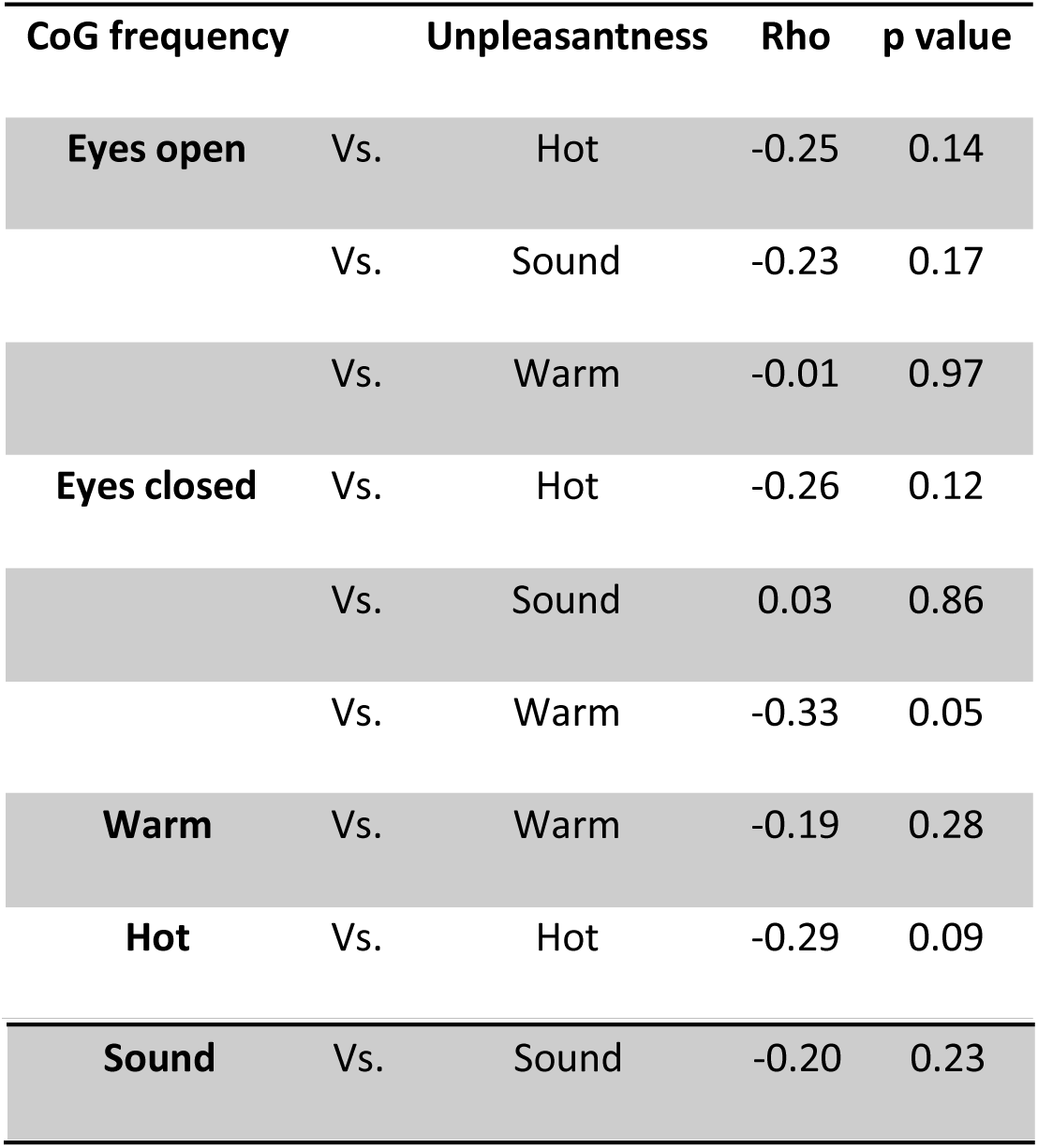
Coefficients of correlation between the 8-10 Hz CoG alpha frequency and unpleasantness ratings.

### Bayesian Correlations

Table 9 summarizes the output of Kendell’s correlations. The analysis of the wide-spectrum CoG alpha frequency (7-13 Hz) revealed moderate evidence in favour of the null hypothesis, thus rejecting the notion of a negative correlation between alpha frequency and the experience of unpleasantness. On the contrary, the analysis of slow CoG alpha frequency (8-10 Hz) revealed mild evidence of negative correlation between unpleasantness ratings and alpha frequency during closed, open eyes and hot stimulation (Fig. 5 left, centre and right).

**Figure 5.**
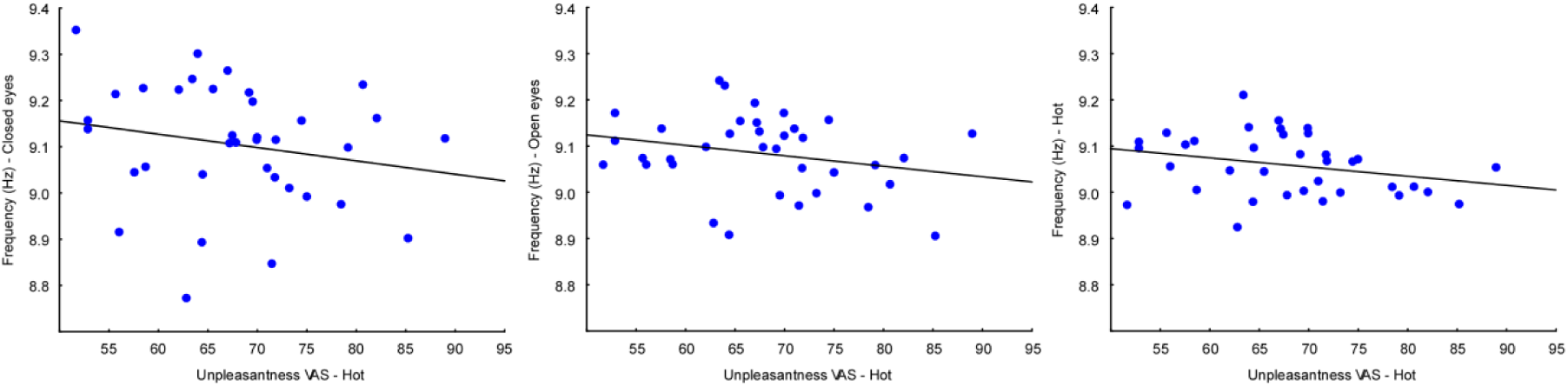
Scatterplots representing the negative correlation between the individual average unpleasantness ratings in the tonic hot condition (x axis) and the 8-10 Hz CoG alpha frequency during closed (left), open eyes (centre) and tonic hot condition (right), respectively. Note the progressive narrowing of the distribution cloud from left to right, thus indicating a strengthening of the correlation in the tonic hot condition relative to the two baselines.

**Table 9.**
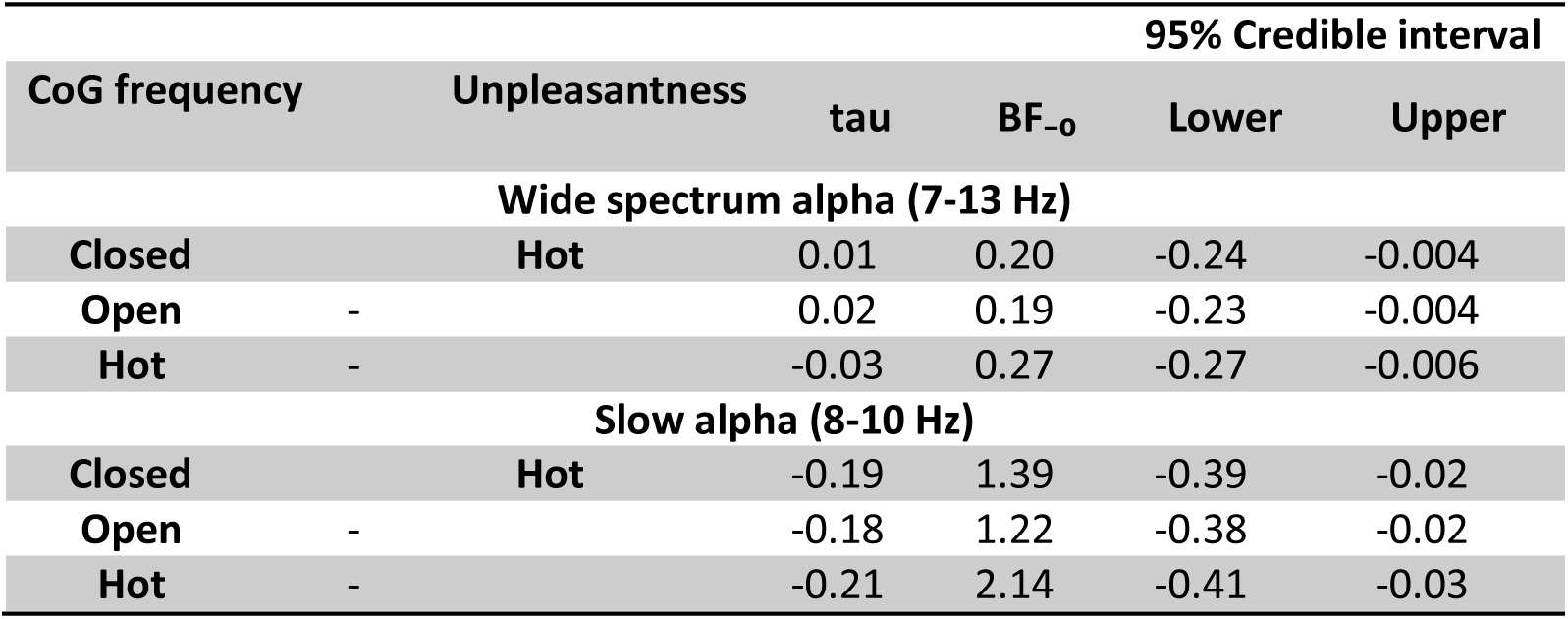
Bayesian correlations between the unpleasantness ratings during hot stimulation and the 8-10 Hz CoG alpha frequency during the closed, open eyes and tonic hot stimulation.

## Discussion

The objective of our study was to test the pain-specificity of EEG alpha oscillations against neutral and perceptually-matched unpleasant non-painful stimulation (Fig 1, right). Our preliminary matching task allowed the participants’ to equalize their perceptual experience by anchoring their sensory evaluation on the unpleasantness of both the hot painful immersion and the distressing prolonged high pitch sound (Fig. 1, left). All the participants were eventually successful in calibrating the two negative affective experiences (Fig. 2). This important methodological feature, together with the introduction of two different types of resting baseline alpha conditions allowed us to quantify the specificity of alpha oscillatory activity (particularly IAF) as an index of prolonged pain stimulation and its relationship with subjects’ perception.

Our findings only partially met the expectations. First, both our analyses demonstrated the expected magnitude reduction of alpha oscillations during prolonged hot water immersion compared with both the baseline recordings and alpha oscillations during warm water immersion (Tables 1 and 5; Figs. 3 and 4). Second, IAF slowed during prolonged hot water immersion compared to baselines only when we restricted the assessment to the lower 8-10 Hz frequency range (Table 7; Fig. 4) whereas IAF accelerated during somatosensory stimulation when we analysed the full 7-13 Hz spectrum (Table 3; Fig. 3). Third, IAP did not correlate with reported unpleasantness nor did IAF (Tables 2, 4, 6, 8). However, when we established an a priori negative correlation between IAF and unpleasantness ratings within a Bayesian analysis framework, we found anecdotal evidence for a negative relationship with the slow alpha range only (Table 9 and Fig. 5). Fourth, our data show no difference in IAF during prolonged hot water immersion compared with an equally unpleasant prolonged auditory experience (Figs. 3 and 4).

The greater IAP during closed and open eyes resting states is a well-established observation (Geller et al., 2014) as well as its suppression during sensory stimulation (e.g. Plöchl, Gaston, Mermagen, König, & Hairston, 2016). In this respect, our finding is no surprise as both phasic and tonic nociceptive painful stimulation are commonly associated with alpha power suppression (Chang, Arendt-Nielsen, & Chen, 2002; Chen & Rappelsberger, 1994; Dowman, Rissacher, & Schuckers, 2008; Giehl, Meyer-Brandis, Kunz, & Lautenbacher, 2014; Peng, Babiloni, Mao, & Hu, 2015).

Our approach to the activation of the nociceptive system was based on a 5 minute immersion in hot water and is comparable to that used by several previous studies (Granot et al., 2008; Lautenbacher, Kunz, & Burkhardt, 2008; Streff, Kuehl, Michaux, & Anton, 2010; Tousignant-Laflamme, Rainville, & Marchand, 2005). Interestingly, our findings seem to be consistent with either the view of increased alpha frequency during tonic pain (Nir et al., 2010) and during an inflammatory model of pain (Furman et al., 2018). In keeping with Nir et al.’s findings (2010) we have detected an increase of 7-13 Hz IAF during the somatosensory stimulation compared with the baselines. And yet, when limiting the IAF extraction to a slower portion of the spectrum (8-10 Hz) we found the opposite pattern: lower frequency during the hot stimulation than the baseline alpha. This is in turn consistent with what found by Furman et al. (2018) with a clinically relevant approach whereby pain was induced by means of a intradermal capsaicin model, triggering prolonged (>15 min) inflammatory pain. In addition, although weakly, our data seem to support the negative relationship between the increase of unpleasantness associated with the prolonged hot immersion and the slowing of alpha oscillations.

Our work relied on the largest sample size (n=36) in a within subject design compared to previous studies on healthy individuals (Chang et al., 2002; Dowman et al., 2008; Furman et al., 2018; Giehl et al., 2014; Li et al., 2016; Nir et al., 2010; Nir, Sinai, Moont, Harari, & Yarnitsky, 2012). Moreover, we based the analytic approach to the extraction of alpha-related information on advanced signal processing while avoiding selective analysis and double dipping (Kriegeskorte, Simmons, Bellgowan, & Baker, 2009). To dissect the relationship between perception and EEG alpha oscillations we based our analytic approach on a twofold strategy. We first preselected the relevant sensory stimulation alpha oscillation on the basis of the criterion that the target activity should not have been observed during the baseline conditions at the beginning and the end of the experimental session (see, Fig.1 and supplementary material). We then submitted our extracted alpha to a whole-alpha frequency range analysis (7-13 Hz) as well as a narrow-band (8-10 Hz) a priori analysis based on previous research evidence (Sarnthein et al., 2006; Vanneste, Song, & De Ridder, 2018). This approach revealed how IAF can behave as a multifaceted index and question whether IAF may behave differently depending on the duration of the pain sensation and on the underlying mechanism involved. In fact, there may be other factors determining which individuals responds to sensory stimulation with increase rather than decrease of the alpha rhythm. For example, future studies may address whether the temporal extent of the sensory stimulation is linked to a different alpha profile in the individual or if similarly unpleasant non-painful experiences are associated with lower alpha frequency during experimental models of clinical pain or in chronic pain individuals. Notwithstanding our large sample size, we should be cautious in extending the significance of our findings to brain activity in both acute and chronic pain as the type of experience induced in the present study (i.e. tonic/prolonged sensation) is relying on distinct physiological mechanisms than those involved in e.g. acute post-operative pain or chronic conditions associated with central sensitization (Gangadharan & Kuner, 2013; Pogatzki-Zahn, Segelcke, & Schug, 2017). Last but not least, the absence of a difference in alpha IAF for equally unpleasant somatosensory and auditory stimulation (and in fact between the hot and warm stimulation too) suggests that IAF may not be specifically reduced during prolonged pain compared to other negative affective bodily states. Nonetheless, the current findings invite more robust and comprehensive research on the role of EEG alpha rhythm during bodily threatening events. Such note of caution is particularly relevant in light of the recent suggestion to manipulate alpha oscillations through non-invasive brain stimulation (Arendsen, Hugh-Jones, & Lloyd, 2018; Hohn, May, & Ploner, 2019) or sensory entrainment (Ecsy, Brown, & Jones, 2018).

In this vein, we recommend that future research would include proper baseline assessment (i.e. pre and post sensory stimulation) as well as non-painful but similarly unpleasant conditions as to quantify the scale and magnitude of the relationship between IAF and the experience of pain. Such methodological posture can also protect us from spurious effects associated with alpha oscillations triggered by various uncontrolled emotional, attentional and memory processes that may influence perceptual processes (Klimesch, 1999; May et al., 2012).

Altogether, our findings provide mild support to the observation that slower alpha frequency (8-10 Hz) may be functionally associated with the most distinctive feature of pain, i.e its unpleasantness. In addition, they provide no support that IAF can dissociate between aversive affective states triggered by different sensory modalities. Crucially, we identified a complex speeding-slowing frequency pattern that emerges on the basis of how the alpha oscillation is analysed. Therefore, we invite caution in interpreting IAF as a brain biomarker of pain sensitivity and underline the impact of methodological and statistical factors in this quest (Mouraux & Iannetti, 2018). Whilst our results partly support previous literature claiming a slowing of alpha frequency as index of pain sensitivity, they also significantly add to the current state of the art by showing that the experimental design and data analysis approach have substantial impact on the direction and robustness of alpha modulations. Further research will need to address methodological aspects that impact on the specificity and sensitivity of IAF as brain marker of perceptual states.

## Acknowledgments

The authors thank Istvan Laszlo Gyimes for assistance with the experimental set-up, Naomi Limbachiya and Irene Gigante for their assistance with data collection. Vincenzo Romei is supported by the BIAL Foundation (Grant 204/18).

## Notes

Conflict of interest: The authors report no conflict of interest.

